# Effects of development and parental care on Hamilton’s force of selection

**DOI:** 10.1101/2022.10.27.513516

**Authors:** Christoph Netz

## Abstract

The force of selection describes the sensitivity of population growth to changes in life history parameters, with a focus usually on the survival probabilities from one age class to the next. Importantly, according to Hamilton the force of selection generally decreases after the onset of reproduction, thereby providing a possible explanation for patterns of senescence. A second characteristic feature is that the force of selection remains constant up to the age of first reproduction. This latter observation, however, rests on the assumption that offspring become independent from their parents right after birth. I show here in a minimal model that if offspring are reliant on their parents, either during early embryonal development or via parental care at later stages, the force of selection on offspring survival generally increases up until the age at which offspring become independent. This provides a possible explanation for the commonly observed pattern of decreasing mortality during early ontogeny. Further, genes acting during recurrent life stages are observed to experience a heightened force of selection compared to genes that act strictly age-specifically, demonstrating the need to develop a mechanistic understanding of gene activation patterns through which to consider life history evolution.

## Introduction

The patterns of mortality observed in natural populations can be understood from an evolutionary perspective either as the product of optimization under constraints or through the balance between selection and mutation. These competing approaches are exemplified by the two major evolutionary theories of ageing, the Antagonistic Pleiotropy theory (Williams 1957) and the Mutation Accumulation theory (Medawar 1952). Under Antagonistic Pleiotropy, patterns of mortality are determined by functional trade-offs between early and later stages of life, whereas Mutation Accumulation considers mortality at a given life stage to be determined by the balance between deleterious mutations and the strength by which such mutations are selected against. Trade-offs and mutation regimes are very difficult to determine empirically, but as long as trade-offs are absent, the force of selection acting on life history parameters can be derived from the observed survival and reproduction rates themselves: Hamilton (1966) defined the force of selection as the differential of the growth rate over the logarithm of survival probability:

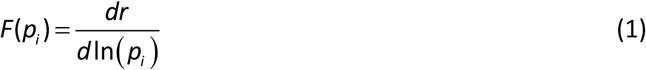

This expression has some interesting properties. First, the force of selection remains constant through the early stages of life before the onset of reproduction, where decreasing cohort sizes are balanced by an increase in prospective reproduction. Second, after reproduction commences, the force of selection is bound to decrease with age. These conclusions however hinge upon the scale, at which variation in survival probabilities occurs (Baudisch 2005). Hamilton assumed that variation occurs relative to the magnitude of survival probabilities, which would for example be the case if the probability to survive from one stage to the next is the product of a number of probabilities to survive different risk factors, and one of these, a risk factor constant across different life stages, is subject to deleterious mutations. If survival probabilities instead vary by a constant amount independently of their magnitude, or relative to mortality rates, the force of selection comes to depend on the age-specific survival parameters and therefore does not need to remain constant during infancy and may also rise at later stages of life (Baudisch 2005). The question at which scale variation typically occurs will likely not be met with a definitive answer, and these measures should therefore be considered as equally relevant. Depending on mutation regime and life history parameters, the force of selection therefore does not need to stay constant before maturity, and also need not always decline after maturity is reached (e.g. negative senescence, Vaupel et al. 2004, Jones & Vaupel 2017).

Hamilton’s force of selection is hence not as generally applicable as once thought. We still may wish to develop an understanding of how selection pressures behave across different life histories, irrespective of the exact parameterization. If survival probabilities are assumed to be age-independent, and therefore constant across different age classes, the two features of constancy before maturity and monotonous decline beyond are retained, regardless of the scale of variation considered or indeed the patterns of fertility observed. The decline of the force of selection may then still serve as a model explanation for widely observed patterns of functional decline and increased mortality with age, even though exceptions may occur. The supposed constant force of selection during infancy however contrasts with extensive evidence that mortality decreases towards maturity (‘ontogenescence’, Levitis 2011, Levitis & Martínez 2013). Age-structured models generally assume offspring to become independent the moment they are born, and thereby ignore dependencies between parents and offspring (but see Lee 2003, Kahn et al. 2015, Roper et al. 2022). Indeed, offspring of many taxa crucially depend on their parents during substantial parts of ontogeny, and parents are often prevented from beginning a new reproductive cycle while taking care of their young. Hamilton himself already suggested a mechanism termed ‘sibling replacement’, by which the force of selection might increase during successive juvenile stages, but he never expressed these ideas in the form of a rigorous model. In 2003, Lee presented a model incorporating energy transfers between family members that demonstrated the potential for the force of selection to increase during successive juvenile stages. Roper et al. (2022), who considered a spatial model with an explicit focus on kin selection, under certain conditions also observed an increasing force of selection at the juvenile stage. Both models incorporate a large amount of mechanistic detail regarding the interactions between family members, and the generality of these findings with respect to the force of selection is presently unclear. In the following, I will develop a much simpler conceptual model that demonstrates the general tendency of the force of selection to increase during dependent juvenile stages.

## The Model

Consider a population, in which offspring develop over consecutive stages *J*_1_ to *J_n_* while being fully dependent on their parent. These stages could be successive gestational stages from fertilization to birth or juvenile stages during which offspring strictly rely on their parent for food and shelter. For simplicity, assume that adults always only cater to a single offspring at a time. We then denote adult reproductive stages *A*_1_ to *A_n_* that run in parallel to the juvenile stages *J*_1_ to *J_n_*. Adults survive from stage *A_i_* to *A*_*i*+1_ with probability *p_i_*, and if the adult dies, so does the offspring. Juveniles survive from *J_i_* to *J*_*i*+1_ with probability *s_i_* and die with probability 1 − *s_i_*. If the offspring dies, the adult returns to stage *A*_1_ and begins a new reproductive cycle. If juveniles pass stage *J_n_*, the juvenile becomes an independent adult and joins the adults at stage *A*_1_ to begin a new cycle of reproduction. Parameters *p_i_* and *s_i_* thus denote adult and juvenile survival. Since juveniles of a given stage are bound to be accompanied by adults of a corresponding stage, we only need to keep track of adult individuals. For *n* = 4 stages, this results in the life cycle shown in figure 1, yielding the following class-structured model 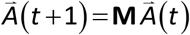:

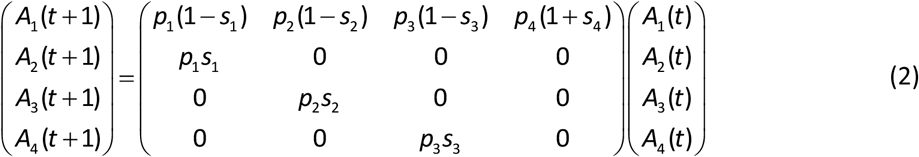

Reproduction is thus finalized only at the end of stage *A*_4_, when juveniles become independent of their parent and join the pool of adults in stage *A*_1_. While the model represents true age classes for juveniles, which can only advance between stages in one direction, for adults the model instead considers different stages of a reproduction cycle that can be undergone multiple times. This also means that for the purposes of this model, vital rates do not depend on parental age.

**Figure 1:**
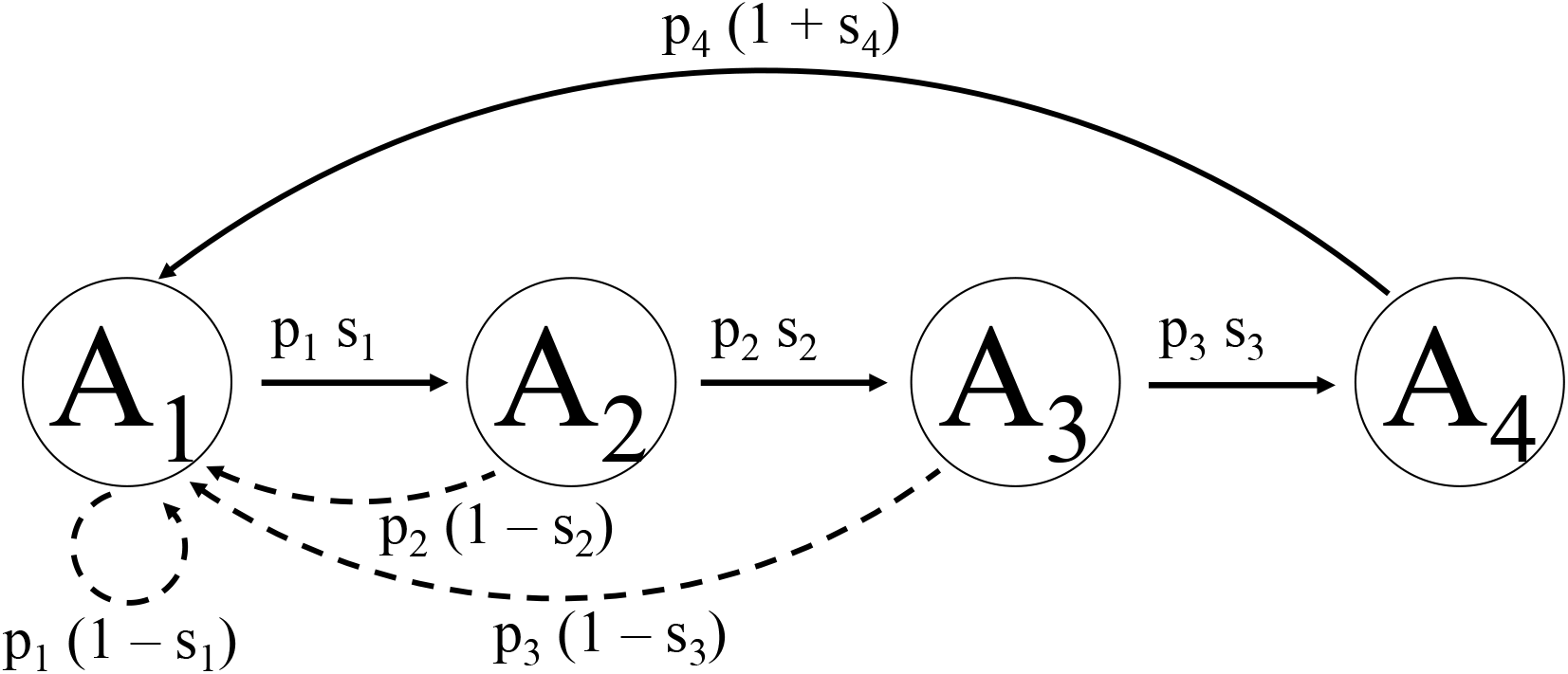
Life cycle with parent-offspring dependence. Adult reproductive stages run from *A*_1_ to *A*_4_, each tracking the number of adults with dependent offspring. *p_i_* and *s_i_* express the survival probabilities of the parent and offspring from one stage to the next. At the end of *A*_4_, offspring join their parents in the pool of mature adults.

A matrix such as the one in eq. 2, that has only non-negative entries, and where all classes are connected, is guaranteed to have a single positive leading eigenvalue *λ*_1_ (Charlesworth 1994). Thus, in the long run, populations will end up growing at a constant rate *λ*_1_, and if *λ*_1_ > 1, such a population grows geometrically. Associated with the leading eigenvalue are a right and left eigenvector. The right eigenvector 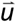 represents the stable class distribution of individuals, which is approached by the population during growth. The left eigenvector 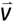 represents reproductive values, indicating the expected future reproduction of an individual in a given stage, relative to the other stages. The force of selection for adult and juvenile survival *p_i_* and *s_i_* may then be calculated using the left and right eigenvectors (see appendix, Caswell 1978, Otto & Day 2011):

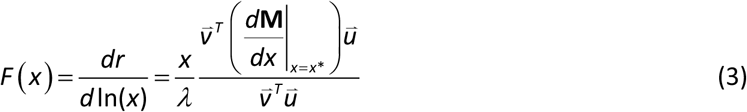

The force of selection on juvenile survival *s_i_* for all stages *i* <*n* then has the pleasingly simple form

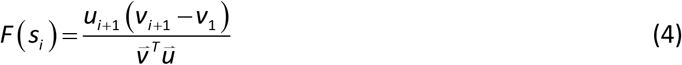

and the force of selection on adult survival *p_i_* for all stages *i* < *n* is

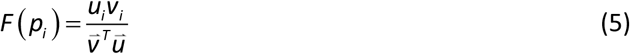

Using the formulas for left and right eigenvectors, 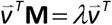 and 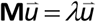 respectively, we first inspect the general behavior of these functions over consecutive life stages, before turning to a specific example where survival rates are the same across all stages.

### General results

The force of selection on the survival of offspring over the first three stages is

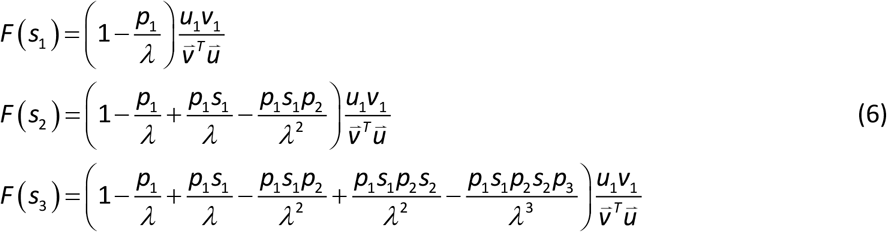

Thus, from stage *i* to stage *i* +1 the force of selection on juvenile survival changes by

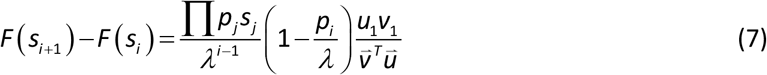

which is positive for all values *p_i_* <*λ*. The force of selection on juvenile survival therefore increases for all biologically realistic values of *p_i_*, as long as population growth is positive.

The force of selection on parental survival over the first three stages is

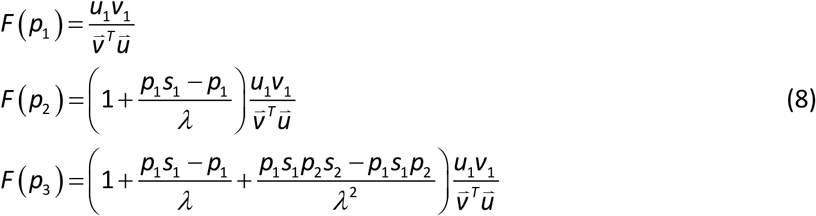

The force of selection on adult survival therefore changes from stage *i* to stage *i* +1 by

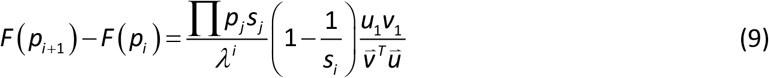

which is equal to or smaller than zero for all biologically realistic values of *s_i_*. The force of selection on adult survival therefore always decreases over successive adult reproductive stages.

We thus observe that when offspring depend on their parents, the force of selection does not stay constant during infancy, but instead always increases. From the beginning of maturity onwards, the force of selection decreases over successive adult stages. Here it should be noted however that the classes represent reproductive stages rather than age classes, which we will turn to at a later point. These results are derived from Hamilton’s classical formula for the force of selection, and hence assume variation in parameters relative to their magnitude.

### Stage-independent adult and juvenile survival rates

In the following, I will present an example in which juvenile and adult survival rates are assumed to be constant across stages. This provides an intuitive understanding for the behavior of the force of selection and allows for the derivation of a general solution. Assuming age-independent juvenile and adult survival rates (*s*_1_ = *s*_2_ = *s*_3_ = *s*_4_; *p*_1_ = *p*_2_ = *p*_3_ = *p*_4_), the force of selection experiences a decelerating increase in the juvenile age classes and a decelerating decrease in the adult reproductive stages that follow (fig. 2, for *s* = *p* = 0.9). Notably, the force of selection on adult survival is much higher than for juvenile survival, because the reproductive stages can reoccur multiple times during the lifetime of an adult.

**Figure 2:**
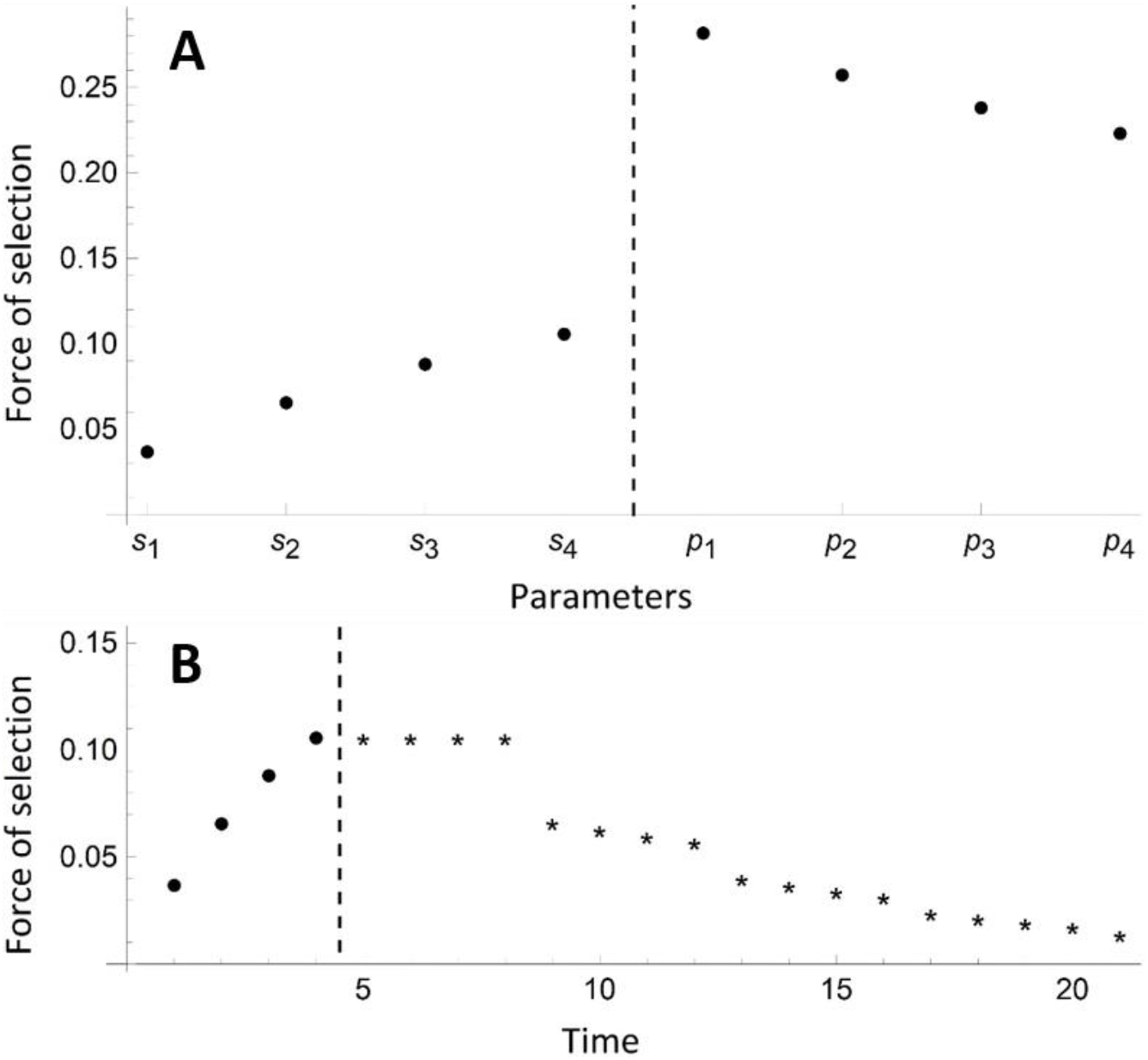
The force of selection during infancy and adulthood. (A) During infancy, the force of selection increases in a slowly saturating manner. The force of selection on adult reproductive stages decreases over consecutive stages, but is much higher than in juvenile age classes. (B) The force of selection on age-specific survival is equal to the force of selection on *s*_1_ to *s*_4_ for the first 4 age classes (•), and from thereon needs to be calculated for adult age classes through eq. 13 (*). Until the first offspring become independent at age 8, the force of selection on adults is equal to the last juvenile stage, and thereafter decreases. The broken line marks the onset of maturity.

The constant juvenile and adult survival rates permit us to derive general expressions for the force of selection on juvenile and adult survival:

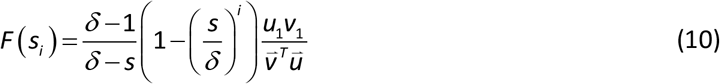

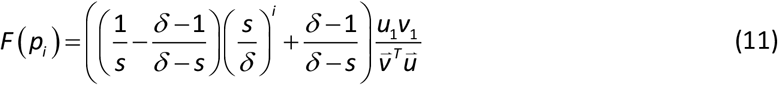

where *δ* is the leading eigenvalue of the matrix **M**/*p* (see appendix). The force of selection on juvenile and adult survival is therefore independent of adult survival *p*, and depends on stage *i* only through the term (*s*/*δ*)^*i*−1^. Since *s*/*δ* is always smaller than 1 (see appendix), this term approaches 0 with increasing *i*. Multiplied with a negative factor in the case of the force of selection on juvenile survival, and with a positive factor in the force of selection on adult survival, the force of selection on juvenile and adult survival are increasing and decreasing saturating functions of *i* (fig. 2).

Following Hamilton, equations 10 and 11 consider the force of selection as the derivative of population growth rate over the logarithm of survival probability. If the derivative is taken over the untransformed survival parameter, mortality rates etc., the force of selection is considered on a different scale and scaling factors that depend on stage-specific survival enter the equation (Baudisch 2005, eq. 5). If parameters are constant across all stages however, as assumed above, this scaling factor is a constant and the overall shape of the derived functions is preserved. While not universal, we can therefore still speak of a general tendency of the force of selection to increase during infancy even if alternatives to Hamilton’s formulation of the force of selection are considered and mutations affect survival probabilities in a variety of ways.

### Force of selection on adult age classes

So far, we considered the force of selection on juvenile and adult survival parameters. Juveniles progress through stages in one direction, and therefore stages represent true age classes. Adult survival however occurs in the context of a reproductive cycle, as shown in figure 1, and the force of selection on adult survival thus relates to reoccurring life stages. The force of selection on adult age classes can then be calculated as follows.

Consider a modified transition matrix **M**° with **M**°_1,4_ = *p*_4_, whereby offspring release at the end of stage 4 is excluded. This matrix can be used to track a single cohort of individuals over different stages and multiple time steps. Adjusting for growth rate, we can calculate the fraction of the stable age distribution due to a cohort of a certain age:

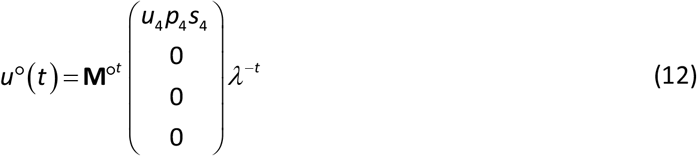

Using the same reproductive values as before, and differentiating over all *p_i_*, we can calculate the force of selection acting on the survival of adults at a given age.

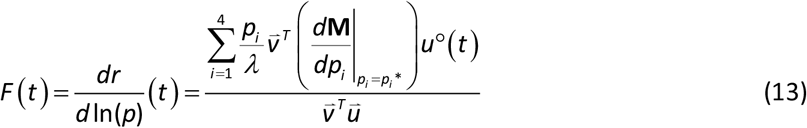

In this case, and in keeping with standard theory, the force of selection remains constant up until the first reproductive cycle is completed at the age of 8, and decreases thereafter (fig. 2, stars).

## Discussion

I have demonstrated here that the selection force generally increases over successive juvenile stages in which offspring depend on their parent. Intuitively, offspring death during early developmental stages is much less costly to parents than during later stages, because the parental investment of time and mortality risk increases over consecutive stages of the reproductive cycle. For example, if offspring die after two time steps, parents lose twice as much time, and experienced twice the mortality risk, than if offspring die after one time step. The mechanism of ‘sibling replacement’ proposed by Hamilton therefore delivers a possible explanation for diminishing mortality rates during early ontogeny. Particularly if matings are easy to come by, the force of selection on offspring survival may thus be initially quite low, and from there increases along with the time/mortality risk invested by parents, until offspring become independent.

Patterns of mortality are determined by the force of selection in mutation-selection balance, or when mutations occur that have opposing fitness effects at different age classes. The model then suggests that mortality rates should decrease from conception until offspring gain independence, remain stable while juveniles are independent but have not begun reproduction, and begin to increase from the time of first reproduction. This seems to be supported by empirical evidence that reports falling mortality rates during ontogeny in a wide range of taxa (‘ontogenescence’, Levitis 2011). In humans, mortality rates continue to fall from birth until the age of 10 to 13 (Barclay 1954, Comfort 1964, Mace 2000).

Further, fetal death rates appear to decrease dramatically over the first trimester of pregnancy, where overall miscarriage risk is reported to drop from 9.4% at 6 weeks of gestation to 4.2% at 7 weeks, 1.5% at 8 weeks, 0.5% at 9 weeks and 0.7% at 10 weeks (Tong et al. 2008). The preclinical rate of pregnancy loss may be yet higher, with 30% of pregnancies after successful implantation lost (as indicated by hCG-testing, Wilcox et al. 1988, Zinaman et al. 1996). These empirical findings can be plausibly explained by our model, but it is important to keep in mind the limitations of a simple measure such as Hamilton’s force of selection; if selection is so strong as to completely deplete any occurring variation, or if drift dominates over selection, the force of selection has no influence on observed rates of mortality. Additionally, if the force of selection is calculated from simple life history data on survival and reproduction, functional trade-offs or evolutionary conflicts that may exist between parents and offspring are ignored. So how much of the observed patterns of ontogenescence could be due to the diminished force of selection in early life?

Extrinsic mortality risks that are common during youth, e.g. due to starvation or predation, may be to a large part unavoidable and not subject to genetic variation or selection. On the other hand, the high rates of failure during the first trimester of pregnancy may easily be under such weak selection that deleterious mutations can accumulate. Medical research indeed seems to indicate a broad variety of causes for early pregnancy failures, e.g. failure of gene activation, chromosomal abnormalities and errors of cytokinesis and karyokinesis (Braude et al. 1991, Michel & Tiu 2007), and such a pattern of heterogeneity is more consistent with mutation accumulation than selection under trade-offs (Austad & Hoffman 2018). If there was a trade-off between mortality in early stages and developmental robustness in later stages, equation [10] would indicate that alleles increasing early-life mortality in favor of developmental robustness later on should be favored. Another important aside is the assumption that genes act age-specifically, the effects of which is observed in the contrast between the force of selection on adult reproductive stages and adult age classes. Developmental stages occur but once in the lifetime of most individual organisms, but reproductive cycles can be undergone multiple times, and this alone increases the force of selection on recurrent or persistently active genes. From a mechanistic point of view, it indeed seems plausible that developmental stages entail a greater number of age-specific genes than age classes during adulthood.

The model I presented here is highly conceptual in nature, and some clear limitations should be highlighted. Individuals have only a single offspring at a time, thus there is no resource competition between siblings (Kahn et al. 2015), and offspring is either dead or alive. The dependency of offspring on their parent is either absolute or non-existent, such that offspring do not optionally rely on their parents for improved survival, as observed in species with facultative parental care (e.g. dung beetles, Capodeanu-Nägler et al. 2016). Such extensions are feasible, but the specification of costs and benefits of parental care becomes nontrivial, with different modelling options to be explored. The models that so far have observed an increasing force of selection before the age of first reproduction were in fact mechanistic models that considered interactions between relatives in greater detail (Lee 2003, Roper et al. 2022). And finally, because the model considers adult reproductive stages rather than age classes, vital rates are independent of parental age. As we have seen, this still permits the calculation of an age-dependent force of selection, but it prevents the incorporation of age-dependent adult survival rates as well as offspring survival rates that depend on the age of the parent (maternal effect senescence, Moorad and Nussey 2016, Ivimey-Cook and Moorad 2020, Hernández et al. 2020). Further work also remains to be done on the force of selection before maturity with regards to potential parent-offspring conflicts (Ronce & Promislow 2010, Kramer et al. 2016) and the effect of density dependence (Mylius & Diekmann 1995, Pen & Weissing 2000, Kokko 2021).

## Acknowledgements

I would like to thank F. J. Weissing and I. Pen for their invaluable advice and comments, as well as the entire MARM group for their constructive feedback. This work was supported by ERC Advanced Grant No. 789240, awarded to F. J. Weissing.

## Appendix

The derivative of the population growth rate *λ* over a parameter *x* is given by the following equation (Caswell 1978, Otto & Day 2011):

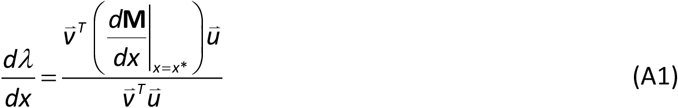

Using the chain rule twice over, and replacing ln*λ* = *r*, we obtain an expression containing the force of selection (eq. 1):

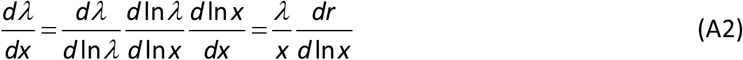

Solving for 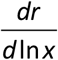 and substituting eq. A1, we obtain eq. 3.

To determine the force of selection acting on juvenile and adult survival at any given life stage, the right and left eigenvectors are used. The elements of the right eigenvector of matrix **M** in equation 3 are given by the following equations:

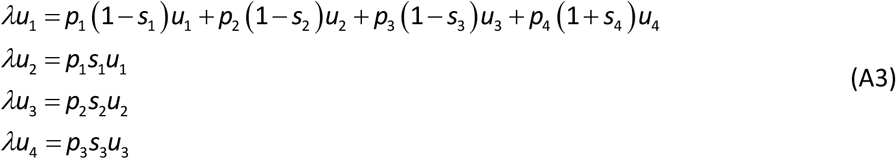

Assuming that *p_i_* and *s_i_* are the same over all stages, we obtain a general expression for the elements of the right eigenvector:

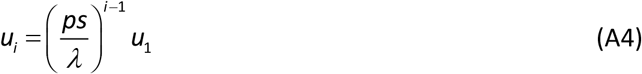

The left eigenvector is determined by the following equations.

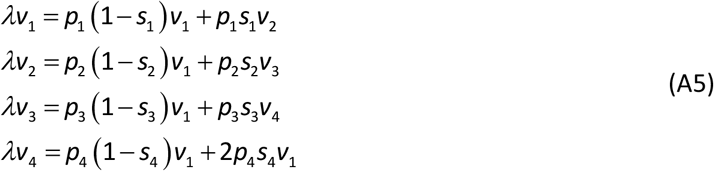

Again assuming that *p_i_* and *s_i_* are the same over all stages, the first term on the right-hand-side is constant. The difference between consecutive elements of the left eigenvector is then

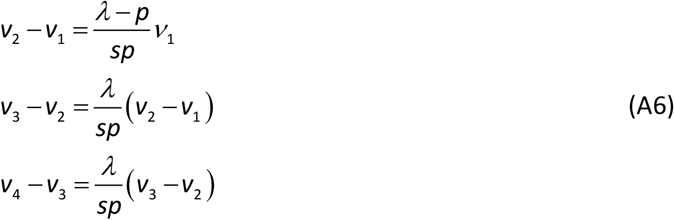

which can be generalized to

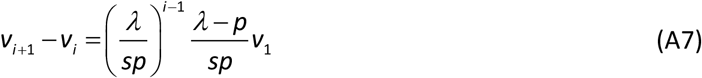

Taking the sum over stages *i* and simplifying, a general expression is obtained for the elements of the left eigenvector:

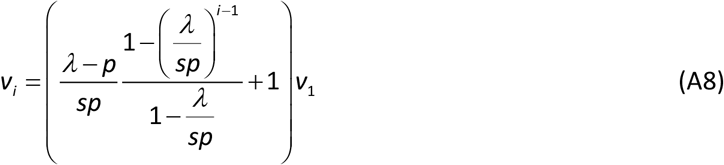

Further, *p* can be factored out of matrix **M**, such that **M** = *p***M**_0_, where **M**_0_ only contains parameter *s*. The leading eigenvalues of **M** and **M**_0_ are then related through *p*: *λ*_1_ = *pδ*_1_, where *δ*_1_ is the leading eigenvalue of **M**_0_, given by the characteristic polynomial:

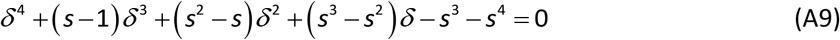

If no offspring survives (*s* = 0), *δ*_1_ is equal to one, and the population is bound to shrink at a rate *λ* = *p*. If *s* >**0**, *δ*_1_ is larger than one and increases with *s*. Plugging A2 and A6 into equations 4 and 5 and replacing *λ* = *pδ*_1_, one obtains the equations 10 and 11 for the force of selection on juvenile and adult survival.

## References

Barclay, G. W. (1954). Colonial development and population in Taiwan. Princeton University Press.

Baudisch, A. (2005). Hamilton’s indicators of the force of selection. Proceedings of the National Academy of Sciences, 102(23), 8263–8268.

Braude, P., Johnson, M., Pickering, S., & Vincent, C. (1991). Mechanisms of early embryonic loss in vivo and in vitro. In The embryo (pp. 1–10). Springer, London.

Capodeanu-Nägler, A., Keppner, E. M., Vogel, H., Ayasse, M., Eggert, A. K., Sakaluk, S. K., & Steiger, S. (2016). From facultative to obligatory parental care: interspecific variation in offspring dependency on post-hatching care in burying beetles. Scientific Reports, 6(1), 1–10.

Caswell, H. (1978). A general formula for the sensitivity of population growth rate to changes in life history parameters. Theoretical population biology, 14(2), 215–230.

Charlesworth, B. (1994). Evolution in age-structured populations (Vol. 2). Cambridge University Press.

Comfort, A. (1964). Ageing. The biology of senescence. Ageing. The biology of senescence. Routledge & Kegan Paul, London.

Hamilton, W. D. (1966). The moulding of senescence by natural selection. Journal of Theoretical Biology, 12(1), 12–45.

Hernández, C. M., van Daalen, S. F., Caswell, H., Neubert, M. G., & Gribble, K. E. (2020). A demographic and evolutionary analysis of maternal effect senescence. Proceedings of the National Academy of Sciences, 117(28), 16431–16437.

Ivimey-Cook, E., & Moorad, J. (2020). The diversity of maternal-age effects upon pre-adult survival across animal species. Proceedings of the Royal Society B, 287(1932), 20200972.

Jones, O. R., & Vaupel, J. W. (2017). Senescence is not inevitable. Biogerontology, 18(6), 965–971.

Kahn, A. T., Jennions, M. D., & Kokko, H. (2015). Sex allocation, juvenile mortality and the costs imposed by offspring on parents and siblings. Journal of Evolutionary Biology, 28(2), 428–437.

Kokko, H. (2021). The stagnation paradox: the ever-improving but (more or less) stationary population fitness. Proceedings of the Royal Society B, 288(1963), 20212145.

Kramer, B. H., van Doorn, G. S., Weissing, F. J., & Pen, I. (2016). Lifespan divergence between social insect castes: challenges and opportunities for evolutionary theories of aging. Current Opinion in Insect Science, 16, 76–80.

Lee, R. D. (2003). Rethinking the evolutionary theory of aging: transfers, not births, shape senescence in social species. Proceedings of the National Academy of Sciences, 100(16), 9637–9642.

Levitis, D. A. (2011). Before senescence: the evolutionary demography of ontogenesis. Proceedings of the Royal Society B: Biological Sciences, 278(1707), 801–809.

Levitis, D. A., & Martínez, D. E. (2013). The two halves of U-shaped mortality. Frontiers in Genetics, 4, 31.

Mace, R. (2000). Evolutionary ecology of human life history. Animal Behaviour, 59(1), 1–10.

Medawar, P. (1952) An Unsolved Problem of Biology. Lewis, London.

Moorad, J. A., & Nussey, D. H. (2016). Evolution of maternal effect senescence. Proceedings of the National Academy of Sciences, 113(2), 362–367.

Mylius, S. D., & Diekmann, O. (1995). On evolutionarily stable life histories, optimization and the need to be specific about density dependence. Oikos, 218–224.

Otto, S. P., & Day, T. (2011). A Biologist’s Guide to Mathematical Modeling in Ecology and Evolution. Princeton University Press.

Pen, I., & Weissing, F. J. (2000). Towards a unified theory of cooperative breeding: the role of ecology and life history re-examined. Proceedings of the Royal Society of London. Series B: Biological Sciences, 267(1460), 2411–2418.

Ronce, O., & Promislow, D. (2010). Kin competition, natal dispersal and the moulding of senescence by natural selection. Proceedings of the Royal Society B: Biological Sciences, 277(1700), 3659–3667.

Roper, M., Green, J. P., Salguero-Gomez, R., & Bonsall, M. B. (2022). Inclusive fitness forces of selection in an age-structured population. bioRxiv, 2022.06.02.494506.

Tong, S., Kaur, A., Walker, S. P., Bryant, V., Onwude, J. L., & Permezel, M. (2008). Miscarriage risk for asymptomatic women after a normal first-trimester prenatal visit. Obstetrics & Gynecology, 111(3), 710–714.

Vaupel, J. W., Baudisch, A., Dölling, M., Roach, D. A., & Gampe, J. (2004). The case for negative senescence. Theoretical Population Biology, 65(4), 339–351.

Wilcox, A. J., Weinberg, C. R., O’Connor, J. F., Baird, D. D., Schlatterer, J. P., Canfield, R. E.,… & Nisula, B. C. (1988). Incidence of early loss of pregnancy. New England Journal of Medicine, 319(4), 189–194.

Williams, G. (1957). Pleiotropy, natural Selection and the evolution of senescence. Evolution, 11(4):398–411.

Zinaman, M. J., Clegg, E. D., Brown, C. C., O’Connor, J., & Selevan, S. G. (1996). Estimates of human fertility and pregnancy loss. Fertility and Sterility, 65(3), 503–509.

